# Foxd1 is required for terminal differentiation of anterior hypothalamic neuronal subtypes

**DOI:** 10.1101/236737

**Authors:** Elizabeth A. Newman, Jun Wan, Jie Wang, Jiang Qian, Seth Blackshaw

## Abstract

Although the hypothalamus functions as a master homeostat for many behaviors, little is known about the transcriptional networks that control its development. To investigate this question, we analyzed mice deficient for the Forkhead domain transcription factor *Foxd1. Foxd1* is selectively expressed in neuroepithelial cells of the prethalamus and hypothalamus prior to the onset of neurogenesis, and is later restricted to neural progenitors of the prethalamus and anterior hypothalamus. During early stages of neurogenesis, we observed that Foxd1-deficient mice showed reduced expression of *Six3* and *Vax1* in anterior hypothalamus, but overall patterning of the prethalamus and hypothalamus is unaffected. After neurogenesis is complete, however, a progressive reduction and eventual loss of expression of molecular markers of the suprachiasmatic, paraventricular and periventricular hypothalamic is observed. These findings demonstrate that *Foxd1* acts in hypothalamic progenitors to allow sustained expression of a subset of genes selectively expressed in mature neurons of the anterior hypothalamus.

**Highlights:**

1. *Foxd1* is broadly expressed in neuroepithelial cells of the hypothalamus and prethalamus.
2. *Foxd1* mutants show severe defects in anterior hypothalamic development, although prethalamic development is only modestly affected.
3. Loss of *Foxd1* results does not affect initial patterning of the hypothalamus, but leads to a progressive loss of expression of markers specific to neurons of the suprachiasmatic, paraventricular and periventricular nuclei.
4. *Foxd1* regulates expression of multiple transcription factors expressed in developing anterior hypothalamus.

## 1 Introduction

The hypothalamus is a master regulator of a broad range of homeostatic processes and innate behaviors, including circadian rhythms, sleep, blood pressure, thirst, feeding and body weight, control of the neuroendocrine system, and a range of sexual, emotional and affiliative behaviors (Bedont et al., 2014; Burbridge et al., 2016; Caron and Richard, 2017; Herrera et al., 2017; Lechan and Toni, 2000; Lee et al., 2012; Liu et al., 2017; Morrison, 2016). Despite its behavioral importance, the development of the hypothalamus is poorly understood, largely because of its considerable anatomic and cellular complexity (Braak & Braak, 1992; Flament-Durand, 1980; Lechan & Toni, 2000). Although recently some progress has been made in identifying extrinsic and intrinsic factors that control hypothalamic patterning and neurogenesis, much remains unknown (Bedont et al., 2015; Burbridge et al., 2016; Xie and Dorsky, 2017).

Large-scale analysis of gene expression in the developing mouse hypothalamus has provided a starting point for examining hypothalamic development, and has identified candidate genes that are expressed in discrete spatial domains of early hypothalamic neuroepithelium (Shimogori et al., 2010). Among these genes is the Forkhead domain transcription factor *Foxd1*. Previously identified as a regulator of kidney and retinal ganglion cell development (Carreres et al., 2011; Hatini et al., 1996), *Foxd1* was observed to be broadly expressed in neural progenitor cells of the prethalamus and anterior and tuberal hypothalamus during early stages of neurogenesis by E11.5 (Figure 1A,B). By E14.5, however, *Foxd1* expression was no longer detected in anterior or tuberal hypothalamus, and became restricted to prethalamic neuroepithelium (Figure 1C,D). Although no hypothalamic defects in *Foxd1* mutants have been previously reported (Carreres et al., 2011; Gumbel et al., 2012; Hatini et al., 1996; Levinson et al., 2005), the spatially and temporally dynamic expression pattern of this gene suggests a potential role in patterning the hypothalamus and/or prethalamus.

**Figure 1:**
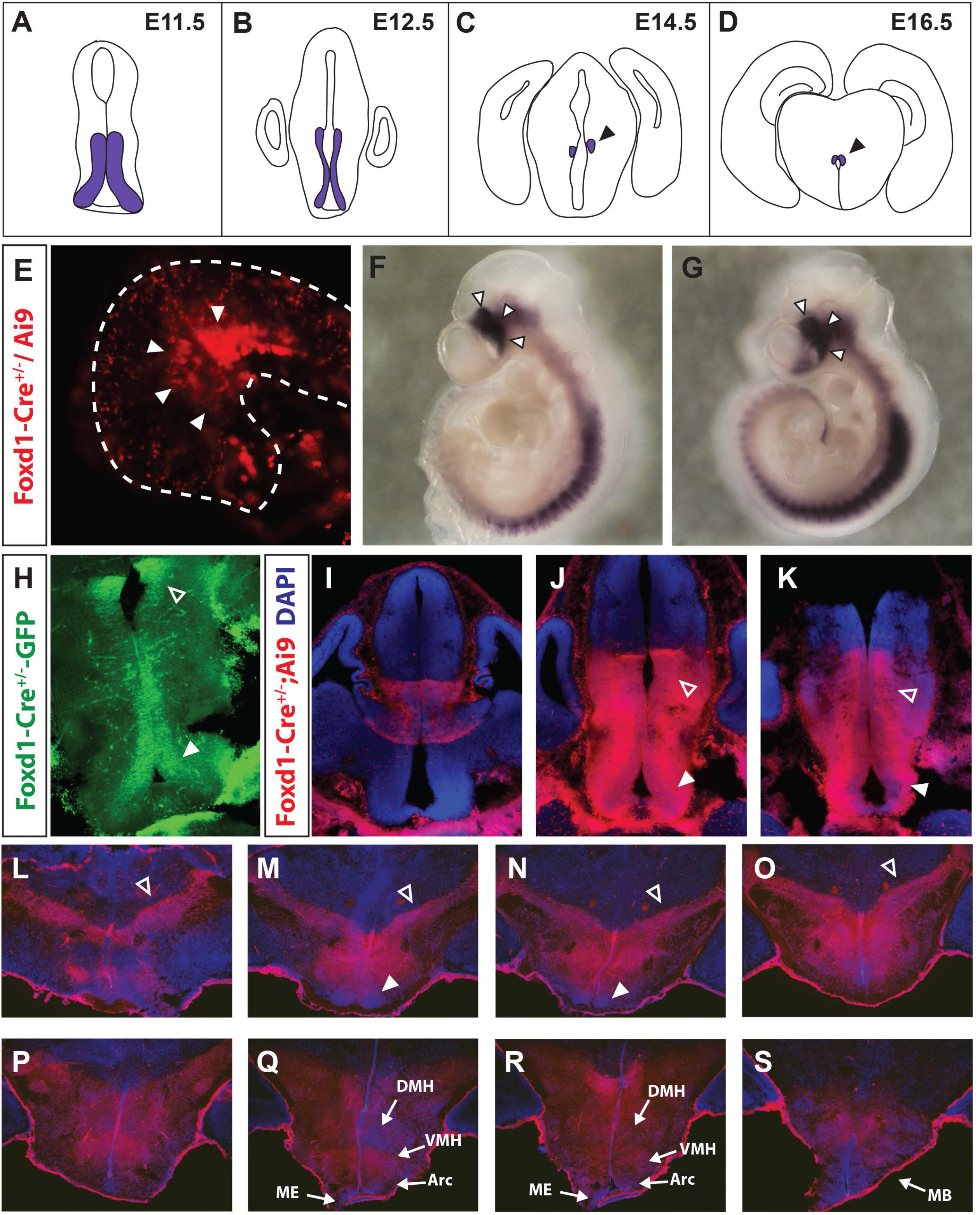
Foxd1 is broadly expressed in hypothalamic and prethalamic neuroepithelial cells. A-D: Schematic showing *Foxd1* mRNA expression in the anterior hypothalamus and prethalamus over the course of development, adapted from Shimogori et al., 2010. E: GFP staining from a *Foxd1-Cre/+; Ai9* E8.5 embryo. White arrowheads indicate the presumptive hypothalamus in the ventral neural tube. F, G: Whole mount *in situ* hybridization of Foxd1 mRNA at E9.0 (F) and E10.0 (G). The hypothalamus is designated by white arrowheads. H: Immunohistochemistry staining against GFP in a *Foxd1-Cre/+* E12.5 brain detects expression of Cre-GFP fusion protein expressed from the *Foxd1* locus. By this age, *Foxd1* is expressed only in the prethalamus (open arrowhead) and the anterior hypothalamus (white arrowhead) along the ventricle. I-K. Despite the limited *Foxd1* expression at E12.5, lineage tracing in E12.5 *Foxd1-Cre/+; Ai9* brains reveals that every cell in the hypothalamus and prethalamus originates from a *Foxd1*-positive lineage. Open arrowheads=prethalamus, white arrowheads=hypothalamus. L-S: DsRed immunohistochemistry against in P0.5 *Foxd1-Cre/+; Ai9* brain reveals a history of Cre activity throughout the prethalamus and hypothalamus. Blue=DAPI. Open arrowheads=prethalamus, white arrowheads=SCN. Sections are arranged in anterior-posterior sequence from top left to bottom right.

In this study, we have investigated the developmental phenotype of *Foxd1* mutants using the region-specific markers identified in our previous work (Shimogori et al., 2010). Despite broad early expression in developing hypothalamus and prethalamus, *Foxd1*-deficient mice show substantially delayed developmental defects that are essentially confined to the anterior hypothalamus. At E12.5, *Foxd1*-deficient mice exhibit only modest developmental defects. Early markers of anterior hypothalamic structures such as the suprachiasmatic nucleus (SCN) are detected at E16.5, but by birth are either lost entirely or substantially downregulated. Moreover, markers of terminal differentiation of the SCN, as well as other anterior hypothalamic structures such as the periventricular nucleus (PeVN) and paraventricular nucleus (PvN) are also lost by birth. The contrast between the selective expression of *Foxd1* in early-stage hypothalamic progenitors, and the delayed appearance of a developmental phenotype, implies that *Foxd1* may control anterior hypothalamic development indirectly by regulating expression of additional transcription factors such as *Six3* and *Vax1*. Alternatively, the delayed developmental phenotype in *Foxd1* mutants may result from altered chromatin accessibility at *cis*-regulatory sequences required for sustained expression of anterior hypothalamic-enriched genes in terminally differentiated neurons. These findings provide new insight into the transcriptional regulatory networks that specify cell identity in the hypothalamus.

## 2 Materials and methods

### 2.1 Mouse breeding and embryo collection

Heterozygous *Foxd1-Cre: GFP* knock-in mice (*Foxd1-Cre/+*)(Humphreys et al., 2008) were mated to each other to generate homozygous animals (*Foxd1-Cre/Foxd1-Cre*) that phenocopy null mutations in *Foxd1*, and are referred to in this manuscript as *Foxd1* mutants. Heterozygous *Foxd1-Cre/+* animals are used as controls for all experiments. Mice were maintained and euthanized according to protocols approved by Johns Hopkins Institutional Animal Care and Use Committee.

### 2.2 *In situ* hybridization

Embryos were post-fixed in 4% paraformaldehyde in 1xPBS, then embedded in gelatin before sectioning. *In situ* hybridization was performed as previously described (Shimogori et al., 2010).

### 2.3 EdU staining and cell counting

Pregnant dams were injected with 50 mg/kg EdU (20 mg/mL in saline) on embryonic day E12.5 and then sacrificed via cervical dislocation 2 hours later. Embryos were post-fixed in 4% PFA, cryopreserved in 30% sucrose, embedded in OCT then cryostat sectioned. EdU staining was performed using the Click-IT EdU imaging kit from Invitrogen, and then the slides were coverslipped using Vectashield. Stained slides were imaged at 40x on a Keyence microscope. Cell counting was conducted using ImageJ on five to six sections per animal. All counting was done blinded and manually.

### 2.4 TUNEL staining

Post-fixed and cryopreserved E12.5 embryos were sectioned and dry mounted. TUNEL staining was performed according to the protocol for cryopreserved tissue in the *In Situ* Cell Death Detection Kit-TMR red (Roche). A positive control was used (3000 U/mL DNAse I). Slides were DAPI stained then coverslipped using Vectashield.

### 2.5 Immunohistochemistry

Embryos were harvested and post-fixed in 4% paraformaldehyde in 1xPBS for a minimum of 16 hours at 4°C. Slides were washed 3x5 min in 1xPBS, then cryopreserved in 30% sucrose in 1xPBS for a minimum of 24 hours at 4°C. Embryos were then embedded in OCT blocks on dry ice, and sectioned on a cryostat at 25μm thickness. Sections were immediately dry mounted onto Superfrost Plus slides and allowed to dry before storing at −20°C. Before beginning the procedure, slides were allowed to dry at room temperature, and then marked with a pap pen to isolate the sections. Slides were incubated in blocking buffer (10% HIHS in 0.1% TX-100 in 1xPBS) for at least two hours at room temperature, before incubating in primary antibody in blocking buffer overnight at 4°C.

Antibodies used are as follows: rabbit anti-GFP 1:1000 (Life Technologies A6455 polyclonal) and rat anti-dsRed 1:1000 (Chromotek 5F8 monoclonal). Slides were washed 3x5 min in 1xPBS-T then incubated with secondary antibody in blocking buffer for two hours at room temperature. Secondary antibodies used were goat anti-rabbit-488 and donkey anti-rat-555. The slides were then washed 5 min in 1xPBS-T before DAPI staining in 1:5000 DAPII in 1xPBS-T for 5 min. Slides were then rinsed 3×5 min in 1x-PBS-T before coverslipping in Vectashield. Stained slides were imaged at 10x on a Keyence microscope.

### 2.6 Whole-mount *in situ* hybridization

All incubations were at room temperature unless otherwise noted, and performed in 24-well mesh plates. E9.0 and E10.0 wild-type embryos were harvested in cold 1xPBS-DEPC. E10 embryos were cut down the midline to allow for probe access. Embryos were fixed in 4% PFA in 1xPBS-DEPC for approximately one week. After fixation, the embryos were washed 2x15 min in PTw (1% Tween-20 in 1xPBS-DEPC) then dehydrated with 25%, 50%, then 75% methanol in PTw for 15 min each, then 100% methanol in PTw 3x15 min. Embryos were stored overnight at −20°C, before rehydration in 75%, 50%, and then 25% methanol in PTw for 15 min each, followed by PTw 2x15 min.

Embryos were then incubated in 6% hydrogen peroxide in PTw for 60 min, followed by PTw wash 2x15 min. Embryos were then incubated 3x30 min in detergent mix (1% Igepal CA-630, 1% SDS, 0.5% Na Deoxycholate, 50mM Tris-HCl pH 8.0, 1mM EDTA, 150 mM NaCl) followed by a 10 min PTw wash. Embryos were digested in 10μg/mL Proteinase K in PTw for 20 min, post-fixed in 0.2% glutaraldehyde, 4% PFA in PTw for 20 min, then washed in PTw 2x10 min. A prehybridization step followed, with room temperature hybridization buffer added to the embryos before incubating at 70°C for one hour. Probe was added directly to the vials (25μl per 5mL hybridization buffer) and hybridized overnight at 70°C. Following hybridization, embryos were incubated in Solution X 4x45 min. At this point, the protocol is identical to the high-quality *in situ* hybridization protocol up to and including adding the antibody. Following antibody incubation, the embryos were washed in 1xTBST 3x10 min, 5x60 min, then overnight. Embryos were washed in NTMT 3x10 min then incubated in an NBT/BCIP color reaction solution at either room temperature or 4°C. Embryos were then washed in 1xTBST to remove the background signal, and the color reaction was stopped in TE.

### 2.7 RNA-Seq analysis

Hypothalamic tissue was dissected at E12.5 and E17.5 as previously described (Shimogori, et al. 2010) and stored −80°C prior to use. A total of 8a *Foxd1-null* and 10 control (either wild-type or *Foxd1-Cre^+/−^*) embryos were harvested at E17.5, which were pooled into two samples for each genotype, for an n=2. In the litter dissected at E12.5 there were 5 embryos of each genotype, of which 3 were E12.5 and 2 were E12.0, which were also pooled into two samples for an n=2. After genotyping, the RNA was extracted using the RNeasy kit (Qiagen). RNA from individuals of each genotype were then pooled and sent for quality control analysis to determine concentration and purity via Bioanalyzer. The E12.5 RNA had significant gDNA contamination, so DNAse digestion using the Qiagen kit was performed prior to Bioanalzyer analysis of isolated RNA. A cDNA library was generated and barcoded for each pooled RNA sample, than sequenced to a paired-end read depth of 100 cycles using Illumina HiSeq 2500. The raw RNA-Seq results were mapped onto mm9 genome (NCBI37) using the default parameters in Tophat and Cufflinks (Trapnell et al., 2012) to generate gene expression FPKM (fragments per kilobase of exon per million). Cuffdiff was used to determine differentially expressed genes between the control and mutant samples. Differentially expressed genes with a q-value of <0.05 were considered significant.

### 2.8 Statistical methods used

For the EdU cell counting, the number of EdU-positive cells was taken as a fraction of the total area of the hypothalamus measured in μm^2^. The values of the sections were averaged for each individual, and then compared using a two-tailed t-test. Two animals were used for each genotype.

## 3. Results

### 3.1 Foxd1 is broadly expressed in neuroepithelial cells of the prethalamus and hypothalamus

We first sought to expand upon our previous study of *Foxd1* expression (Shimogori et al., 2010) by more fully exploring the spatial and temporal expression pattern of *Foxd1* in the developing hypothalamus and prethalamus. For these studies, we used a mouse line in which a Cre-GFP fusion had been inserted in frame with the start codon of the endogenous *Foxd1* locus (Hatini et al., 1996; Humphreys et al., 2008), which we then crossed to the Cre-dependent tdTomato reporter line *Ai9* (Madisen et al., 2010). Using this approach, we observed Cre activity as early as E8.5 in the region of the presumptive hypothalamus (Figure 1E). Using whole-mount *in situ* hybridization, *Foxd1* mRNA was found to be strongly expressed in prethalamic and hypothalamic neuroepithelium at E9.5 (Figure 1F) and E10.5 (Figure 1G), as well as in paraxial mesoderm, as previously described (Hatini et al., 1994). Using immunohistochemistry against GFP in heterozygous *Foxd1-Cre-GFP* knock-in animals, we found that *Foxd1* expression was restricted to hypothalamic and prethalamic neural progenitors at E12.5 (Figure 1H), consistent with our previous *in situ* hybridization analysis (Shimogori et al., 2010). Analysis of *Foxd1-Cre-GFP; Ai9* mice at E12.5, however, revealed that virtually the entirety of both hypothalamic and prethalamic regions expressed tdTomato, demonstrating that these cells arise from Foxd1-expressing progenitors (Figure 1I-K). This was the also seen in *Foxd1-Cre-GFP; Ai9* mice at P0.5 (Figure 1L-S), consistent with previous analysis of adult *Foxd1-Cre-GFP; Ai9* mice (Salvatierra et al., 2014).

### 3.2 *Foxd1*-deficient mice show modest reductions in *Six3* and *Vax1* expression at E12.5, but no changes in cell proliferation or death

We generated *Foxd1*-deficient mice by creating homozygous *Foxd1-Cre-GFP/ Foxd1-Cre-GFP* mice, which, as previously reported, show no detectable expression of native protein and die neonatal (Hatini et al., 1996). We first analyzed these mice at E12.5, conducting section *in situ* hybridization using a panel of markers that label discrete regions within prethalamus and/or hypothalamus (Figure 2A) (Shimogori et al., 2010). Most probes tested, including *Isl1, Rax, Sim1, Arx, Lhx1* and *Lhx5* (Figure 2B–M), showed no detectable difference in either expression pattern or signal intensity. In addition, no changes in either proliferation or cell death were observed at this age in *Foxd1*-deficient embryos (Figure 3).

**Figure 2:**
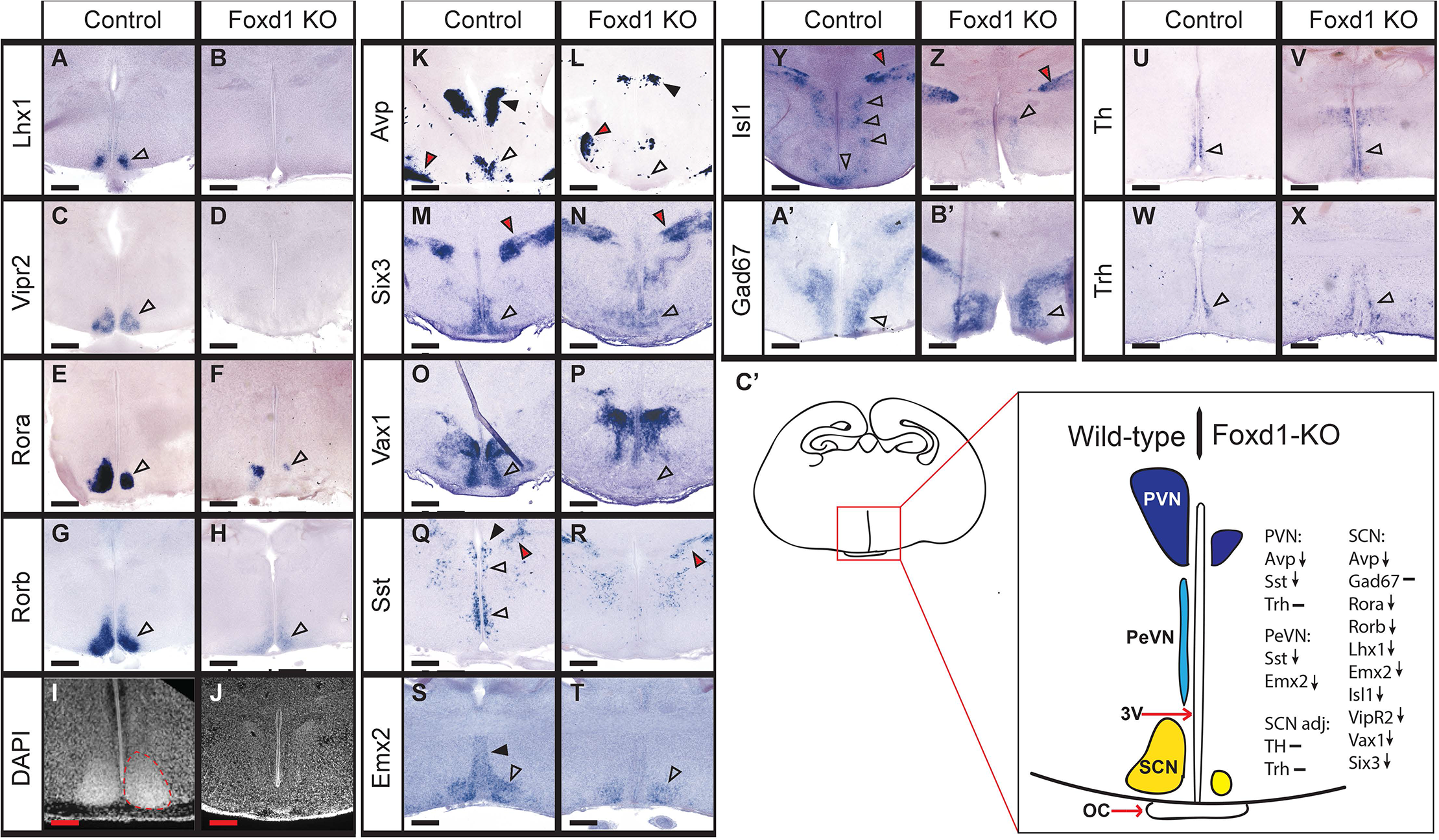
At E12.5, most hypothalamic markers are unaffected by loss of Foxd1. A: Coronal schematic demonstrating the organization of the hypothalamus and prethalamus shown in B-Q. B-Q: *In situ* hybridization on E12.5 heterozygote control and *Foxd1* mutant coronal sections. Most selective markers of the ventral AH (open arrowheads, BE, H-K), and prethalamus (black arrowheads, B, C, H-K) were unaffected in *Foxd1* mutant brains. However, *Vax1* and *Six3* expression was reduced in the ventral AH (open arrowheads, F, G, N, O) and *Six3* expression in the prethalamus was also reduced (black arrowheads indicate the prethalamus, red arrowhead shows missing expression domain, F, G). *Sim1* expression in the presumptive PVN was unaffected (black arrowheads, L, M).

**Figure 3:**
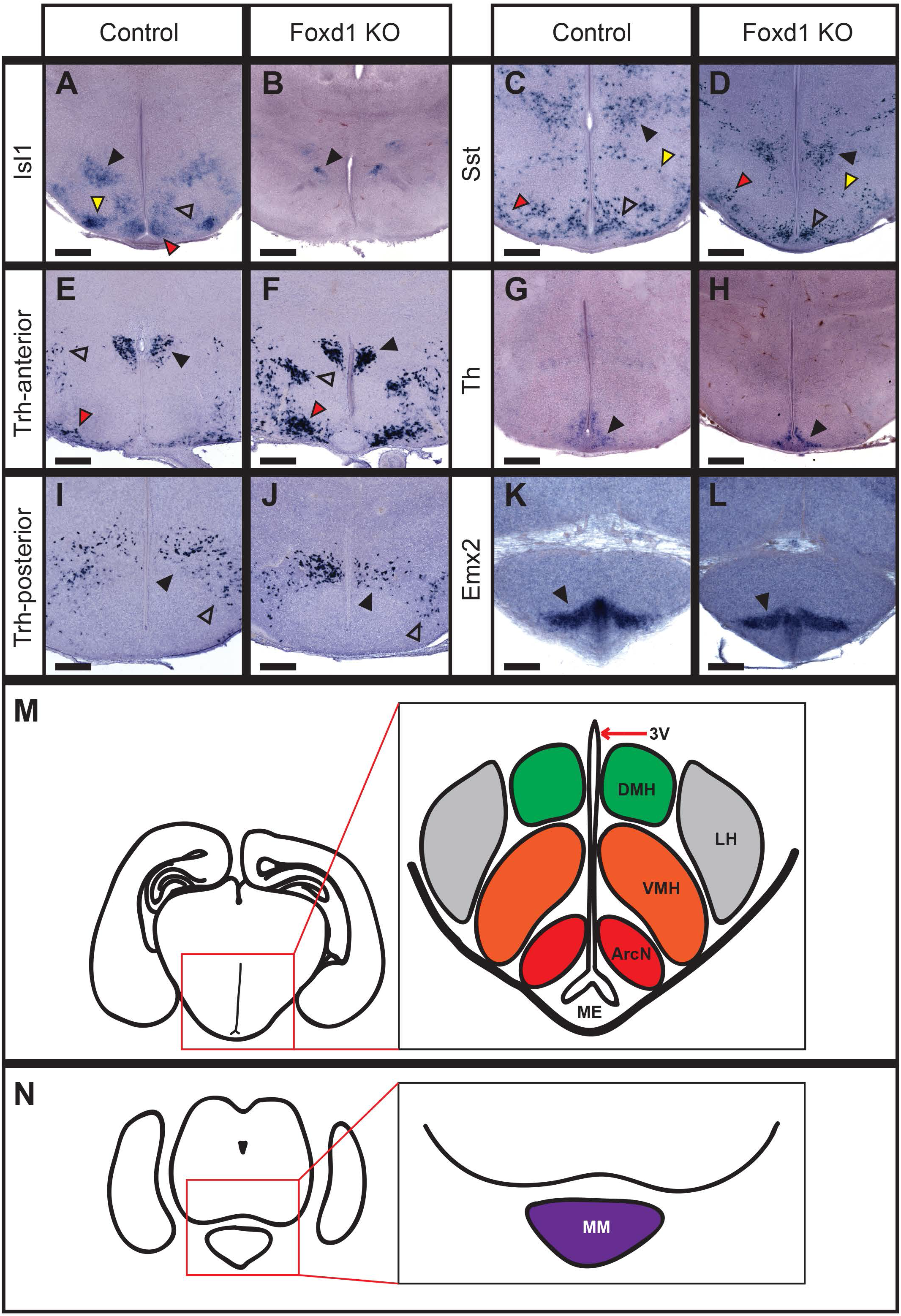
Proliferation and apoptosis are not affected by loss of Foxd1. A-B: Immunohistochemistry staining against EdU (red) and DAPI (blue) on E12.5 heterozygote control and *Foxd1* mutant coronal sections after a 2hr EdU pulse. Red scroll bars indicate the counted portion of the section containing the prethalamus and hypothalamus. C: Quantification of EdU-positive cells relative to area (μm^2^) in control versus *Foxd1* mutant (n=2). D-E: There was no obvious change in the number of TUNEL-positive cells in *Foxd1* mutants.

In contrast, expression levels of the transcription factors *Six3* (Figure 2N,O) and *Vax1* (Figure 2P,Q) showed reduced expression in the ventral hypothalamus of *Foxd1*-deficient embryos. *Six3* also showed reduced expression in its prethalamic expression domain, with a more dramatic reduction observed in the ventricular zone (Figure 2O; red arrowhead) than in the mantle zone (Figure 2O; black arrowhead).

### 3.3 *Foxd1*-deficient mice show reduced expression of a subset of anterior hypothalamic markers by E16.5

At E16.5, we observed selective defects in the expression of molecular markers for individual neuronal populations in the SCN and prethalamus in *Foxd1*-deficient embryos (Figure 4A). Although the broadly-expressed gene *Isl1* did not show altered expression (Figure 4B,C), we observed a reduction in the expression levels of the SCN markers *Rorb* (Figure 4D,E) and *Lhx1* (Figure 4F,G). Expression of somatostatin (Sst) was also disrupted in the thalamic reticular nucleus – a prethalamic structure – but was not detectably altered in lateral hypothalamus (Figure 4H,I).

**Figure 4:**
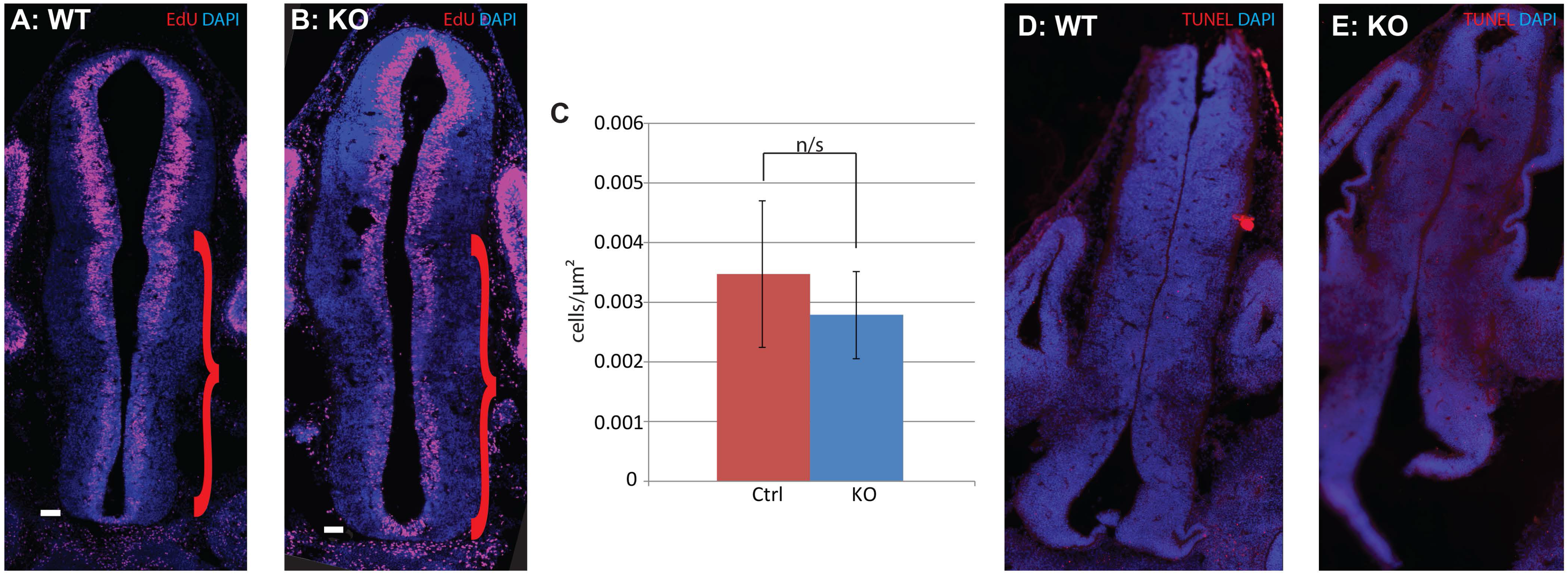
A subset of anterior hypothalamic markers show reduced expression at E16.5. I: A schematic demonstrating the organization of the hypothalamus and prethalamus at the coronal plane shown in A-I. A-I: *In situ* hybridization on E16.5 heterozygote controls and *Foxd1* mutant brains. B-E: A subset of SCN-specific markers show modestly reduced expression at E16.5 (black arrowheads in B-E indicate the SCN). F, G: At E16.5, *Sst* expression is not yet detectable in the PeVN (black arrowheads), but shows altered organization in the prethalamus (open arrowheads). H, I: *Isl1*, which is broadly expressed in anterior hypothalamus, was unaffected at E16.5 (black arrowheads).

### 3.4 *Foxd1*-deficient mice show SCN agenesis and severe reduction of a subset of anterior hypothalamic markers by P0.5

By P0.5, severe defects in SCN, PeVN and PVN differentiation were observed in *Foxd1*-deficient mice. Expression of *Lhx1*, a master regulator of terminal SCN differentiation (Bedont et al., 2014; Bedont and Blackshaw, 2015; Bedont, et al. 2017), was completely absent (Figure 5A, B), as was expression of *Vipr2*, an essential mediator of circadian entrainment in the SCN (Figure 5C,D) (Harmar et al., 2002). *Rora* and *Rorb*, two selective markers of the SCN and key transcriptional regulators of the core circadian clock, showed substantially reduced expression (Figure 5E–H). The histology of the SCN was also severely disrupted, with DAPI staining showing no remaining detectable nuclear organization (Figure 5I, J).

**Figure 5:**
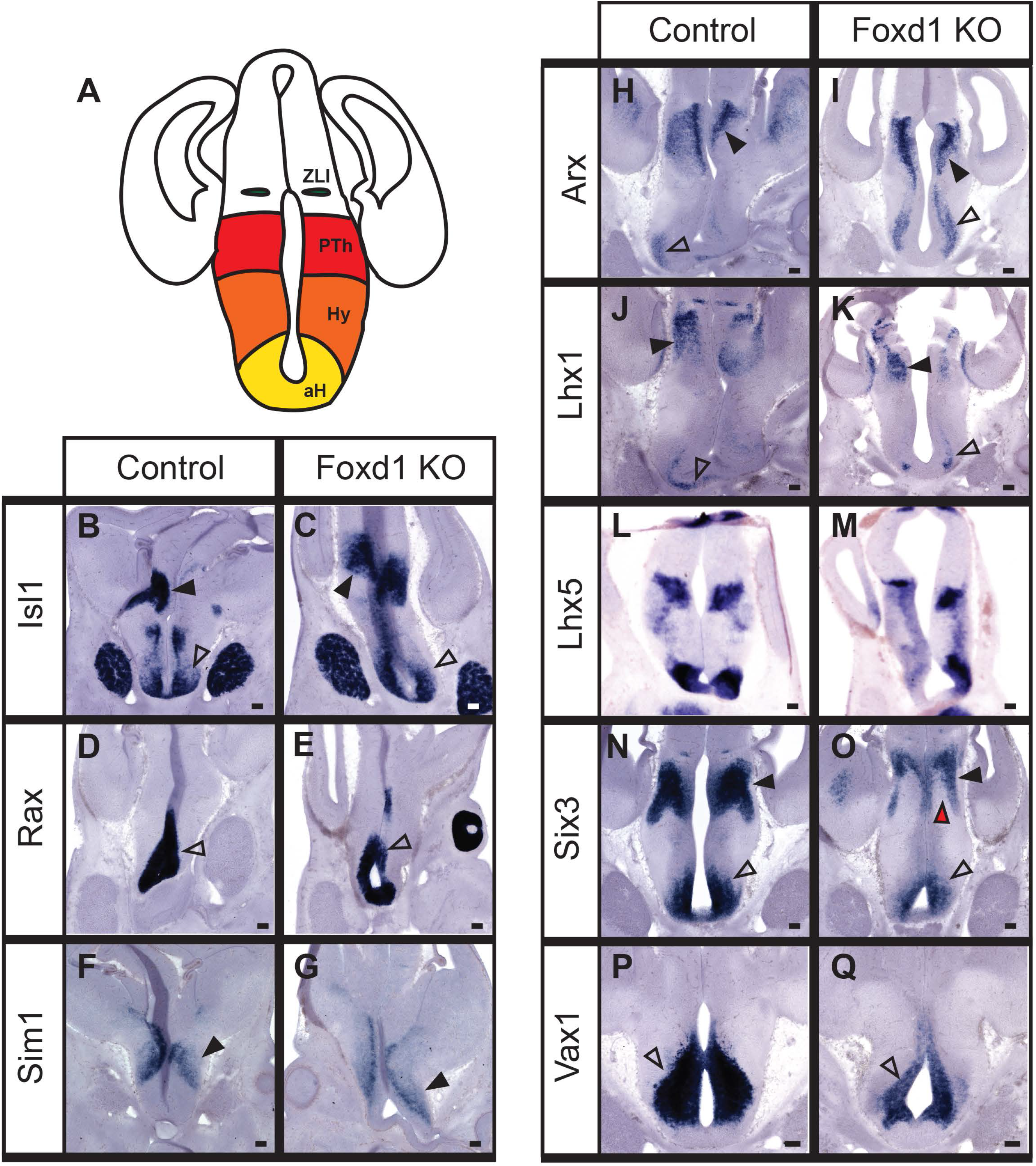
Loss of *Foxd1* results in a broad and selective loss of anterior hypothalamic markers at P0. A-H; K-B’: *In situ* hybridization of coronal sections from P0 heterozygote controls and *Foxd1* mutant brain. Expression of selective markers of the SCN was either absent or greatly reduced in *Foxd1* mutants relative to controls (open arrowheads, A-H, K-P, S, T). *Gad67* expression, however, was unaffected (open arrowheads, W, X), as were *Th* and *Trh* (open arrowheads, Y-B’), which are expressed in regions of the AH adjacent to the SCN. In sharp contrast to wild-type animals (red dashed outline, I) the majority of *Foxd1* mutants showed no evidence of an organized SCN (J). Markers of the PVN were also negatively affected by the loss of *Foxd1* (black arrowheads, K, L, Q, R), as was *Sst* in the PeVN (open arrowheads, Q, R). The broadly expressed hypothalamic marker *Isl1* was strongly reduced in anterior hypothalamus, but not in prethalamus (open arrowheads, U, V). Other prethalamic markers were unaffected (red arrowheads, M, N, Q, R). C’: Schematic summarizing developmental defects in the anterior hypothalamus. Expression of anterior hypothalamic markers was either reduced (down arrow) or unaffected (−).

Genes that are expressed in SCN, as well as other anterior hypothalamic regions, also showed altered expression. *Avp* expression in SCN was virtually absent, and was substantially reduced in PVN, although its expression in SON was unaffected (Figure 5K, L). Expression of *Six3*, an essential transcriptional regulator of SCN development (VanDunk et al., 2011), showed dramatically reduced expression in the SCN region, as did the transcription factor *Vax1* (Figure 4M–P). *Sst* expression was absent from the PeVN and PVN (Figure 5Q, R), and expression of the transcription factor *Emx2* was likewise absent from the PeVN, but continued to show expression in the subparaventricular zone (vSPZ) (Figure 5S, T). *Isl1*, which is broadly expressed in the prethalamic-derived reticular nucleus, as well as the anterior and tuberal hypothalamus at this age, also showed substantially reduced expression throughout the hypothalamus, although expression in reticular nucleus was unaffected (Figure 5U, V; Figure 6A, B).

**Figure 6:**
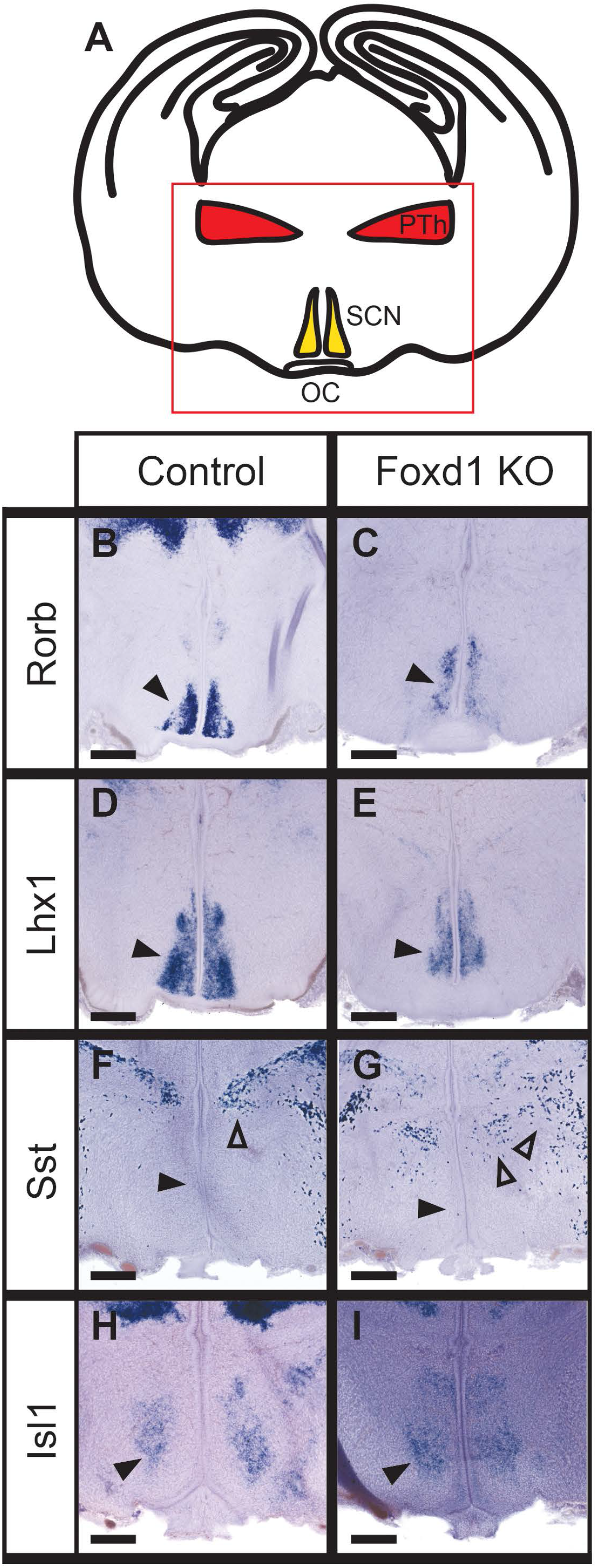
Posterior and lateral hypothalamic structures are largely unaffected in Foxd1-deficient animals at P0. A-L: *In situ* hybridization on P0 heterozygote control and *Foxd1* mutant coronal sections. Expression of *Sst* (open arrowheads, C, D), and *Th* (black arrowheads, G, H) in the ArcN were unchanged in *Foxd1* mutants, but Isl1 expression (red arrowheads, A, B) was lost. DMH markers were unaffected (black arrowheads, C, D, I, J), with the exception of *Isl1* (open arrowheads, A, B), which was also reduced in the ventrolateral VMH (yellow arrowheads, A, B) and LH (black arrowheads, A, B). The lateral hypothalamus had normal *Trh* expression (open arrowheads, E, F, I, J). Markers of the tuberomammillary terminal (red arrowheads, C, D), and the supraoptic nucleus (red arrowheads, E, F), and mammillary nucleus (black arrowheads, K, L) were also unaffected. M, N: Schematics showing the overall organization of posterior hypothalamic nuclei on two different coronal planes. M corresponds to the coronal plane featured in A-D, G-J and N corresponds to sections K, L.

**Figure 7:**
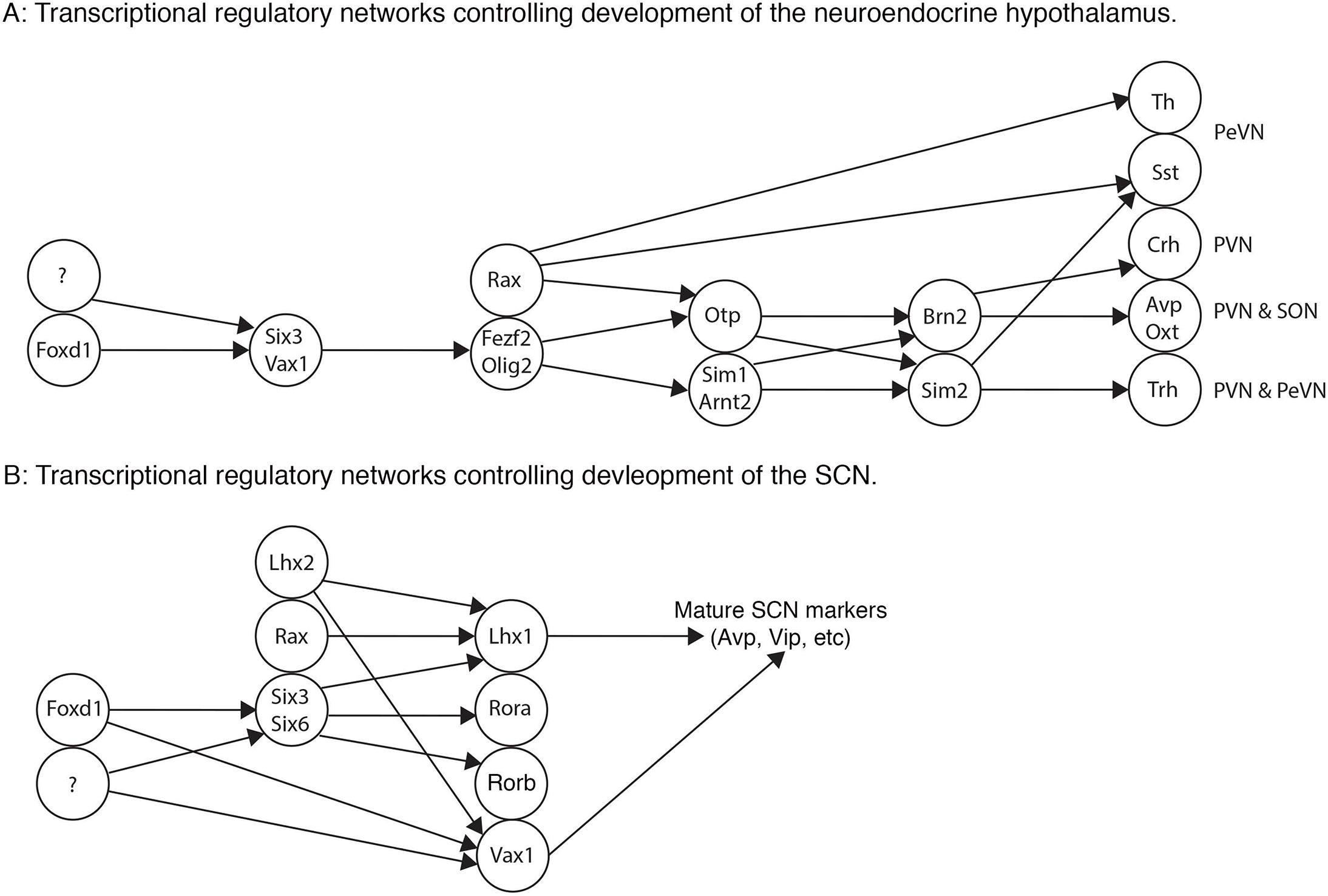
Schematic demonstrating model of the role of Foxd1 in anterior hypothalamic development. A: The known transcriptional regulatory pathways governing development of the dorsal anterior hypothalamus. We hypothesize that *Foxd1* regulates *Six3* and *Vax1*, which in turn may regulate *Fezf2* and *Olig2*. B: The transcription factors known to be involved in development of the SCN. We hypothesize that *Foxd1* is controlling SCN development through regulation of *Six3* and *Vax1*. In both the dorsal anterior hypothalamus and the SCN, we hypothesize that *Foxd1* is working in conjunction with other, unknown transcription factors, due to the incomplete loss of *Six3* and *Vax1* in the *Foxd1-null* animals.

Not all anterior hypothalamic markers showed reduced expression in *Foxd1*-deficient mice at P0.5, however. Expression of *Th* in A14 dopaminergic cells of the PeVN was unaffected (Figure 5Y, Z), as was *Trh* expression in the PeVN (Figure 5A’, B’) and the PVN (Figure 6E, F). *Gad67*, which is expressed in GABAergic neurons throughout the anterior hypothalamus (O’Hara et al., 1995), likewise was not affected in *Foxd1*-deficient mice (Figure 5W, X). Despite the fact that reduced expression of *Six3* was observed in prethalamus at E12.5 (Figure 2N, O), its expression in thalamic reticular nucleus was unaffected at P0.5 (Figure 5N, red arrow). Likewise, no changes in *Sst* and *Isl1* expression were seen in the prethalamus (Figure 5T, V red arrowhead), despite the fact that hypothalamic expression of both genes was severely disrupted, and the fact that prethalamic *Sst* expression was disrupted at E16.5 (Figure 4G). Finally, aside from decreased *Isl1* expression, no changes in expression of markers of the tuberal and posterior hypothalamus were observed (Figure 6E–N).

### 3.5 RNA-Seq analysis identified additional genes showing altered expression in *Foxd1*-deficient mice at E12.5 and E17.5

To further characterize changes in gene expression resulting from loss of function of *Foxd1*, we conducted RNA-Seq analysis of E12.5 and E17.5 hypothalamus and prethalamus derived from heterozygote controls and *Foxd1*-deficient mice. At both ages we observed a dramatic downregulation of *Foxd1* mRNA levels, as expected, but otherwise saw essentially no other overlap in genes that showed significantly altered expression at E12.5 and E17.5 (q<0.05). We observed 2 genes that were upregulated and 49 genes that were downregulated at E12.5, and 16 genes that were upregulated and 10 that were downregulated at E17.5. Several of the genes upregulated at E17.5 are selectively expressed in telencephalic structures, such as *Foxg1, Neurod2* and *Tbr1* (Table 2), which suggest that these might reflect contamination of these samples by telencephalic structures.

At E12.5, we also observed substantially reduced expression of *Zfhx2* and *Zfhx3*, two transcription factors selectively expressed in anterior hypothalamus, as well as *Six3* (Table 1). Other transcriptional regulators that show both reduced expression in *Foxd1* – deficient mice and selective expression in embryonic anterior hypothalamus include the transcription factor *Tox2*, the histone demethylase *Kdm6b*, and the transcriptional coregulator *Rere* (Eichele and Diez-Roux, 2011; Lein et al., 2007).

**Table 1:**
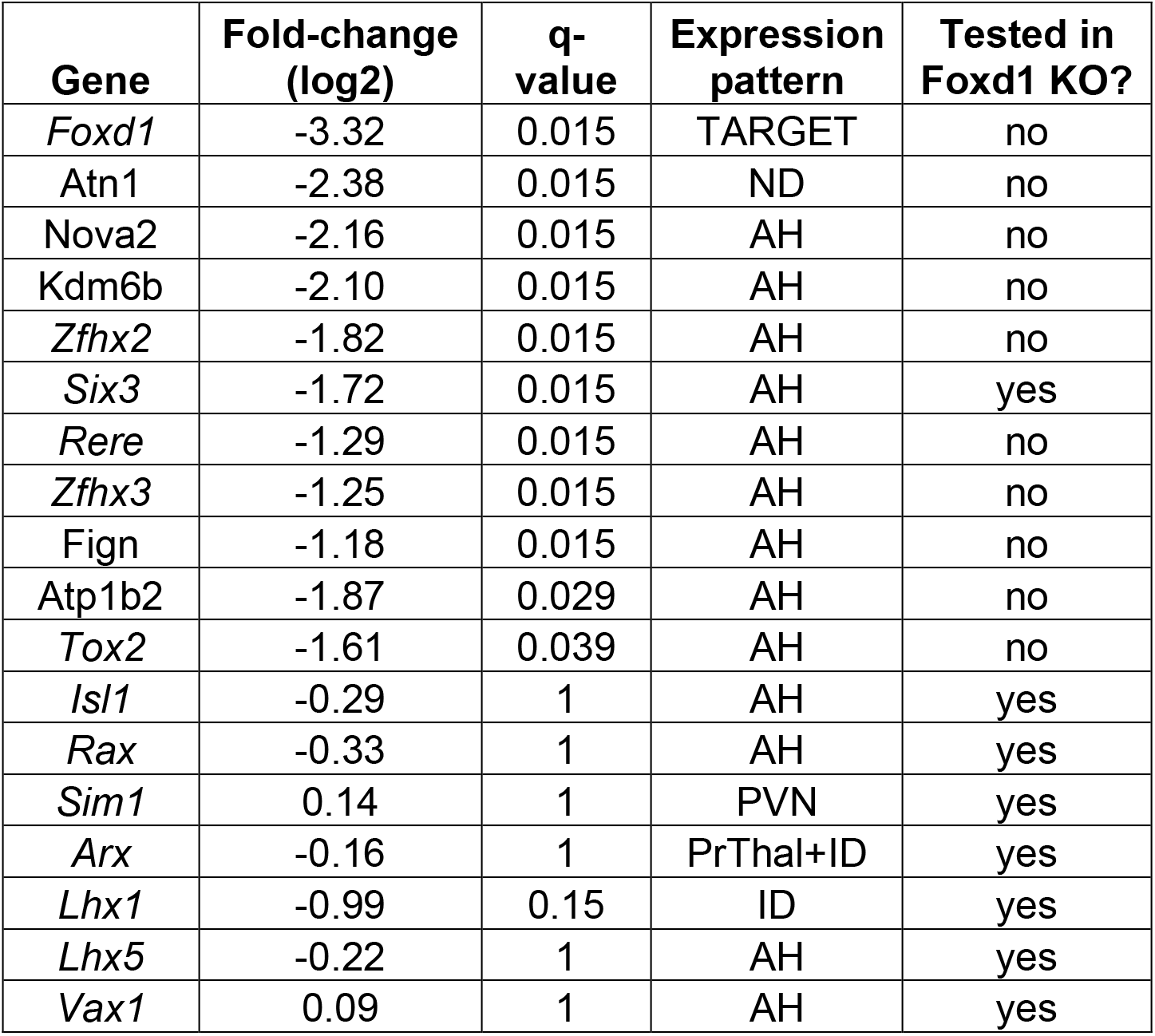
RNA-Seq analysis of control and *Foxd1*-deficient E12 hypothalamus. RNA-Seq analysis of E12 hypothalamus from heterozygote control and *Foxd1* mutants. Genes are listed that show significant differential expression and/or were analyzed by i*n situ* hybridization. Two replicates from E12 and E12.5 were combined for this analysis. Differentially expressed genes were determined by q-value<0.05. Transcription factors and coregulators are highlighted in italic.

With the exception of *Pou4f1* and *Irx1*, which likely represent dissection contaminants from habenula, all of the genes that are significantly downregulated in E17.5 *Foxd1* – deficient mice are selectively expressed in anterior hypothalamus. These include the neurohormones *Avp, Oxt* and *Crh;* the transcription factor *Vax1*, and the cell adhesion molecule *Cdh23* (Table 2).

**Table 2:**
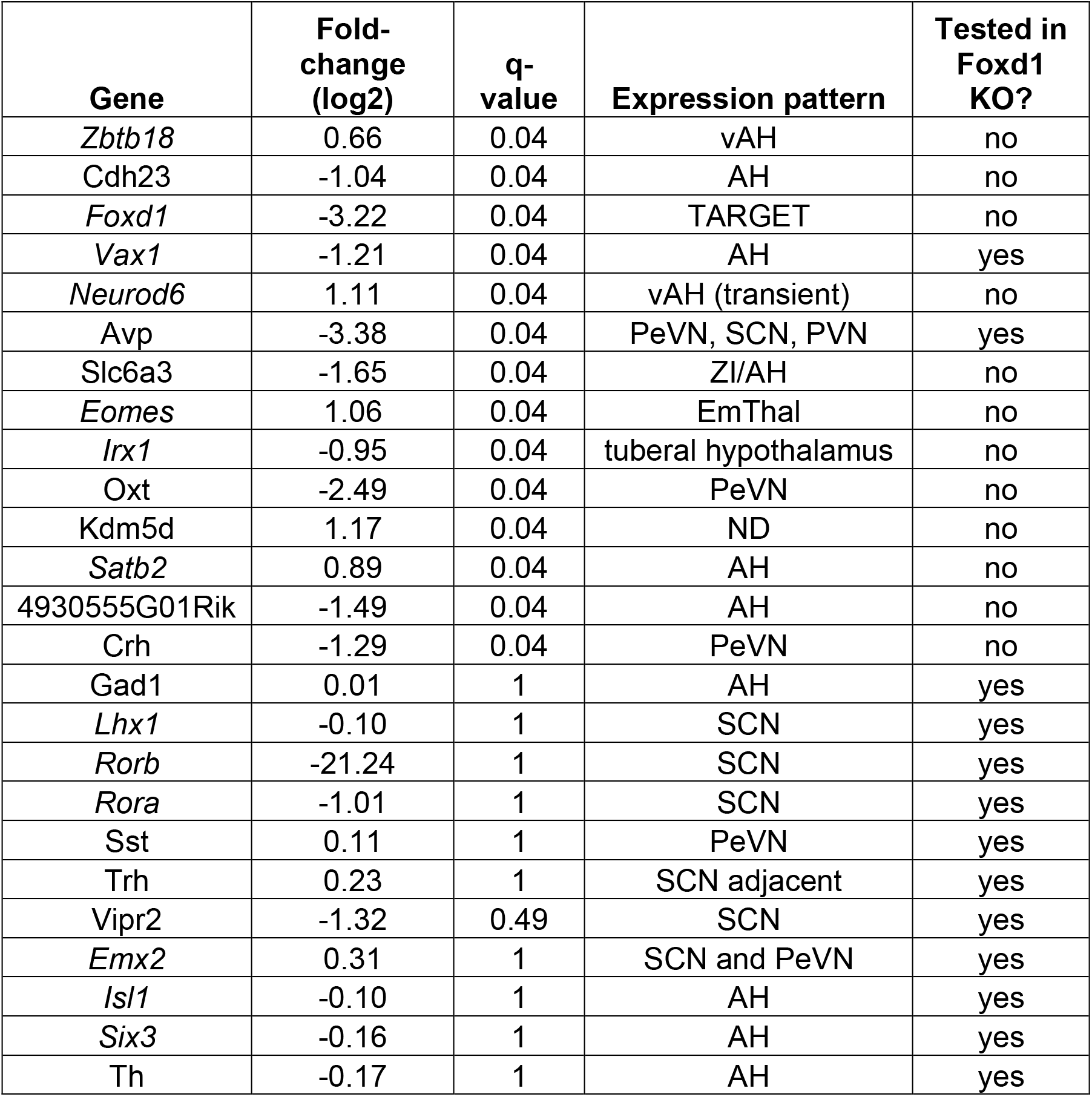
RNA-Seq analysis of control and *Foxd1*-deficient E17,5 hypothalamus. RNA-Seq analysis of E17.5 hypothalamus from heterozygote control and *Foxd1* mutants. Genes are listed that show significant differential expression and/or were analyzed by i*n situ* hybridization. Differentially expressed genes were determined by q-value<0.05. Transcription factors and coregulators are highlighted in italic.

## 4. Discussion

In this study, we have identified *Foxd1* as an essential regulator of anterior hypothalamic development. Despite showing strong and selective expression in early hypothalamic and prethalamic neuroepithelial and neural progenitor cells, the effects of loss of function of *Foxd1* are initially quite modest at E12.5, and are primarily confined to downregulation of a handful of genes that are selectively expressed in developing anterior hypothalamus, including the transcription factors *Six3, Vax1* and *Zfhx3*. Most anterior hypothalamic markers, and all tuberal and posterior hypothalamic markers tested were unaffected at this age, and only *Six3* showed reduced expression in neural progenitors of the prethalamus. At E16.5, when neurogenesis is essentially complete in hypothalamus and prethalamus and *Foxd1* expression is barely detectable, we observed reduced expression of early markers of the developing SCN, including *Lhx1* and *Rorb*, along with reduced *Sst* expression in thalamic reticular nucleus (which is derived from prethalamus). By P0.5, expression of SCN-specific markers is either absent or greatly reduced, and nucleogenesis of the SCN does not occur. In addition, there is a reduction or absence of expression of a subset of PeVN and PVN-specific markers, including *Avp* and *Sst*. Although there appears to be disrupted expression of a limited number of prethalamic markers at earlier timepoints, this is no longer observed at P0.5.

These findings lead us to draw several conclusions about the mechanism by which *Foxd1* regulates the development of the prethalamus and hypothalamus. First, despite its initially broad expression throughout both structures, and selective expression in anterior hypothalamus following the onset of neurogenesis, *Foxd1* is not required for spatial patterning or neurogenesis in either region; no obvious transformations in the identity of individual cell types are observed following *Foxd1* loss of function.

Second, the effects of *Foxd1* loss of function in hypothalamus primarily reflect a failure to either maintain expression of early-onset cell specific genes or to undergo terminal differentiation. An example of the former is *Lhx1*, the earliest detectable selective marker of the developing SCN, which initiates expression normally, but which shows reduced and then absent expression at later stages of development. An example of the latter is *Sst* expression in the PeVN and PVN, which is never initiated. It is, however, likely that failure of terminal differentiation results from disrupted maintenance of expression of master transcriptional regulators, such as *Lhx1* itself in the case of the SCN. It is also important to note that in no hypothalamic region examined was there a complete failure of differentiation. Although the SCN was the most severely affected area, cells in this region continued to express the GABAergic marker *Gad67*, and also showed reduced but still detectable expression of *Rora* and *Rorb*. In the PeVN, *Sst* and *Emx2* expression were lost but *Th* and *Trh* expression were preserved. Likewise, *Avp* and *Sst* expression in the PVN were sharply downregulated but *Trh* expression was unaffected. Taken together, these data imply that *Foxd1* does not regulate cell identity *per se*, but is instead necessary to maintain expression of a subset of cell type-specific genes in anterior hypothalamus.

Third, despite strong and persistent expression of *Foxd1* in prethalamic neuroepithelium, only modest and transient changes in prethalamic markers were observed. Although reduced *Six3* and *Sst* expression were observed in embryonic prethalamus, at P0.5 there were no changes in either the pattern or intensity of prethalamic markers detected. This implies that while *Foxd1* may regulate the kinetics of differentiation of certain prethalamic cell types, it is dispensable for their terminal differentiation.

Finally, the discrepancy between the neural progenitor-specific expression pattern of *Foxd1* and the late onset of the developmental phenotypes observed in *Foxd1* loss-of-function mutants implies that *Foxd1* acts indirectly to control expression of genes associated with terminal differentiation. The most obvious mechanism by which this could occur is by regulation of expression of other transcription factors, which in turn go on to guide terminal differentiation of specific neuronal subtypes.

Several potential candidate factors were identified in our analysis, including *Six3*, *Zfhx3* and *Vax1. Six3* is expressed broadly in the developing anterior hypothalamus, and is one of the most strongly downregulated genes in *Foxd1* mutants, as measured by both RNA-Seq and *in situ* hybridization. Previous work has shown that *Six3* is essential for terminal differentiation of the SCN (VanDunk et al., 2011). The same study also suggested a possible role in regulating *Avp* expression in the PVN (VanDunk et al., 2011). Another gene known to regulate SCN-specific gene expression is the zinc homeodomain factor *Zfhx3*, which is significantly downregulated in RNA-Seq analysis of E12.5 *Foxd1* mutant hypothalamus. Hypomorphic mutations of *Zfhx3* have been shown to lead to reduced expression of SCN-specific genes and altered circadian function (Balzani et al., 2016; Parsons et al., 2015; Wilcox et al., 2017). A final candidate gene is *Vax1*, also shown to be strongly downregulated as measured by both RNA-Seq and *in situ* hybridization, which is also expressed in ventral hypothalamus, and has been reported to be required for expression of SCN-specific genes and maintenance of normal circadian activity rhythms (Hoffmann, et al. 2016).

*Foxd1*-dependent regulation of *Six3, Zfhx3* and *Vax1* may thus account for the severe defects in SCN development observed in *Foxd1* mutants. It is important to note that expression of these genes is reduced but not eliminated in *Foxd1* mutants, implying that additional, as yet unidentified, transcription factors are required to maintain expression of these genes in anterior hypothalamus. Further and more detailed analysis of mice mutant for these genes will be needed to determine whether disruption of these genes also accounts for observed defects in PeVN and PVN.

A second, and not mutually exclusive, potential mechanism of action of *Foxd1* might be to prime the activation of cis-regulatory elements that control expression of cell type-specific genes during late stages of differentiation in hypothalamic neural progenitor cells. Other members of the Forkhead family such as *Foxa1* are known to possess pioneering activity, or the ability to directly bind to closed chromatin and induce a shift to an open conformation (Lupien et al., 2008; Taube et al., 2010). It is thus possible that *Foxd1* selectively targets enhancer elements that are necessary for transcription of critical transcriptional regulators such as *Lhx1* in terminally differentiated neurons, altering their chromatin conformation and rendering them accessible for binding by other cell type-specific transcription factors. Global analysis of chromatin conformation using techniques such as ATAC-Seq (Buenrostro et al., 2013) in *Foxd1*-deficient hypothalamic progenitors can potentially address this question.

## Acknowledgements

We thank W. Yap for comments on the manuscript. This work was supported by NIH R01DK108230, and a Hopkins Synergy Award (to S.B.)

## Author contributions

EAN conducted experiments and analyzed all data. JW and JQ analyzed RNA-Seq data. EAN and SB interpreted data, and wrote the manuscript, with input from all authors.

## Author information

RNA-Seq data is available in GEO. The authors declare no competing financial interests.

